# regLM: Designing realistic regulatory DNA with autoregressive language models

**DOI:** 10.1101/2024.02.14.580373

**Authors:** Avantika Lal, David Garfield, Tommaso Biancalani, Gokcen Eraslan

## Abstract

Cis-regulatory elements (CREs), such as promoters and enhancers, are DNA sequences that regulate the expression of genes. The activity of a CRE is influenced by the order, composition and spacing of sequence motifs that bind to proteins called transcription factors (TFs). Synthetic CREs with specific properties are needed for biomanufacturing as well as for many therapeutic applications including cell and gene therapy.

Here, we present regLM, a framework to design synthetic CREs with desired properties, such as high, low or cell type-specific activity, using autoregressive language models in conjunction with supervised sequence-to-function models. We used our framework to design synthetic yeast promoters and cell type-specific human enhancers. We demonstrate that the synthetic CREs generated by our approach are not only predicted to have the desired functionality but also contain biological features similar to experimentally validated CREs. regLM thus facilitates the design of realistic regulatory DNA elements while providing insights into the cis-regulatory code.

## 1 Introduction

Cis-regulatory elements (CREs), such as promoters and enhancers, are DNA sequences that regulate gene expression. Their activity is influenced by the presence, order, and spacing of sequence motifs [23] that bind to proteins called transcription factors (TFs), similarly to how words and phrases define the meaning of a sentence. Synthetic CREs with specific properties are needed for biomanufacturing as well as numerous therapeutic applications including cell and gene therapy; for example, to maximize activity of a therapeutic gene in the target cell type.

Such CREs are often designed manually based on prior knowledge [9]. Recent studies have used directed evolution [19, 21] and gradient-based approaches [17, 14, 10] for CRE design, in which supervised ‘oracle’ models are trained to predict the activity of a CRE from its sequence, and are then used to edit sequences iteratively until the desired prediction is achieved. However, such approaches are not truly generative and do not necessarily learn the overall sequence distribution of the desired CREs. Instead they may only optimize specific features that have high predictive value. Consequently, the resulting CREs may be out-of-distribution and unrealistic, leading to unpredictable behavior when they are experimentally tested in a cell.

Autoregressive language models, such as Generative Pre-trained Transformer (GPT) can produce realistic content in natural languages [5]. Here, we present regLM, a framework to design synthetic CREs with desired properties, such as high, low or cell type-specific activity, using autoregressive language models in conjunction with supervised models. Although masked language models have been used to embed or classify DNA sequences [13, 7, 2, 8, 24], to our knowledge this is the first time language modeling has been used for DNA in a generative setting.

## 2 Results

### 2.1 regLM adapts the HyenaDNA framework for CRE generation

Several transformer-based foundation models for DNA have been developed [13, 7, 2, 8, 24]. However, these methods are based on masked language modeling which is difficult to use for sequence generation. In contrast, the recent Hye-naDNA foundation models [15] are single-nucleotide resolution autoregressive models trained on the human genome, and are hence suitable for regulatory element design. These models are based on the Hyena operator [16], which uses implicit convolutions to scale sub-quadratically with sequence length.

regLM builds on the HyenaDNA [15] framework to perform generative modeling of CREs with desired properties using prompt tokens. This takes advantage of the resolution and computational efficiency of the HyenaDNA model. Further, the ability to fine-tune pre-trained models which have already learned regulatory features enables even design tasks which lack sufficient labeled data for training.

Given a dataset of DNA sequences labeled with their measured activity (Fig. 1A), we encode the label in a sequence of categorical tokens (‘prompt tokens’), which is prefixed to the beginning of the DNA sequence (Fig. 1B). We train or fine-tune a HyenaDNA model to take the processed sequences and perform next token prediction beginning with the prompt tokens (Fig. 1C). This formulation allows us to use any prior knowledge on sequences in the model explicitly. Once trained, the language model can be prompted with the sequence of tokens representing any desired function. The model, now conditioned on the prompt tokens, generates a DNA sequence one nucleotide at a time (Fig. 1D). In parallel, we train a supervised sequence-to-activity regression model on the same dataset (Fig. 1E), and apply it to the generated sequences to select those that best match the desired activity (Fig. 1F). This combined approach allows us to use the regression model as an oracle like previous model-guided approaches, while the language model ensures that the generated sequences have realistic content. Finally, we provide several approaches to evaluate the generated sequences as well as the model itself (Fig. 1G).

**Fig. 1.**
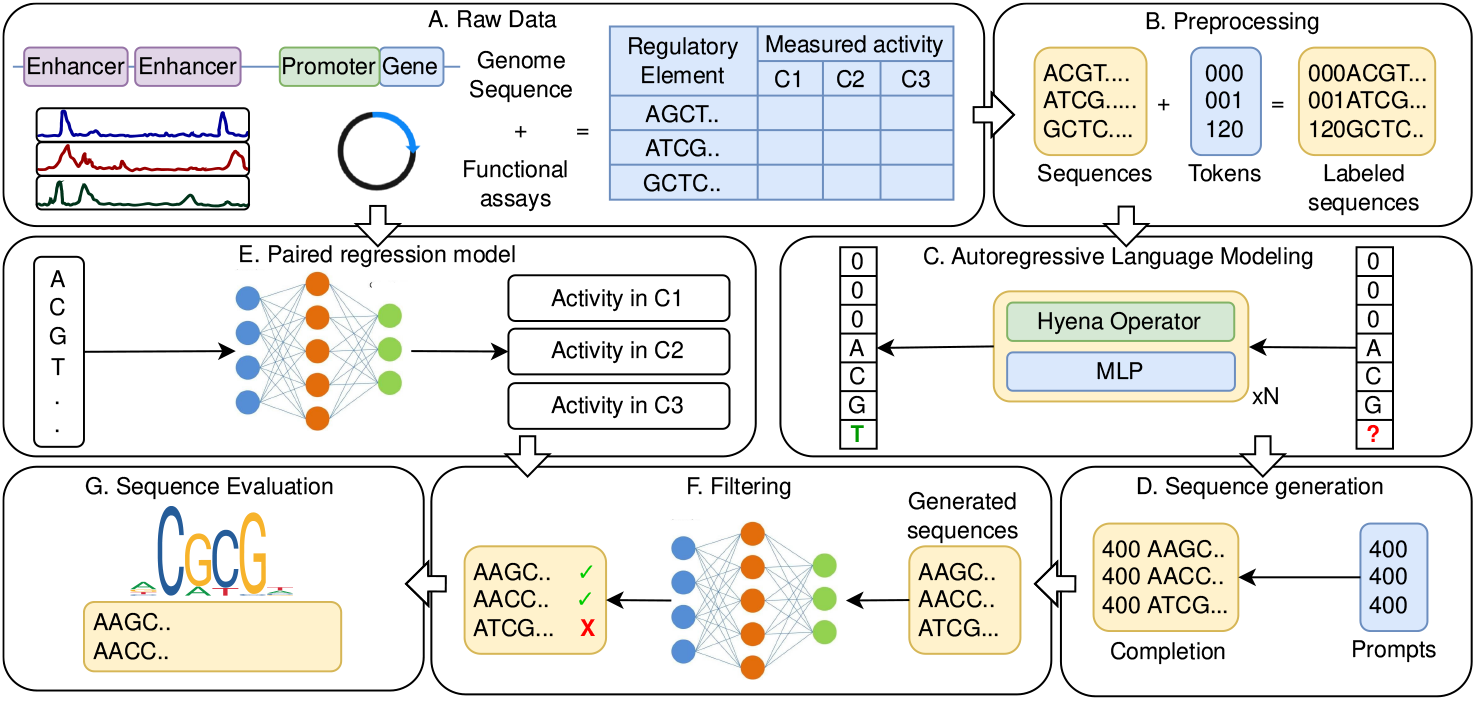
Schematic of regLM. A, B) DNA sequences are prefixed with a sequence of prompt tokens representing functional labels. C) A HyenaDNA model is trained or fine-tuned to perform next token prediction on the labeled sequences. D) The trained model is prompted with a sequence of prompt tokens to generate sequences with desired properties. E, F) A sequence-to-function regression model trained on the same dataset is used to check and filter the generated sequences. G) The regulatory content of generated sequences is evaluated.

### 2.2 regLM generates yeast promoters of varying strength

#### Training and evaluating a regLM model on yeast promoter sequences

We applied the regLM framework to a dataset of randomly generated 80 base pair (bp) DNA sequences and their measured promoter activities in yeast grown in complex and defined media [4, 21]. We prefixed each sequence with a two-token label, wherein each token ranges from 0 to 4 and represents the promoter activity in one of the media (Fig. S1). For example, the label 00 indicates that the sequence has low activity in both media, while 04 indicates low activity in complex medium and high activity in defined medium (Fig. 2A).

**Fig. 2.**
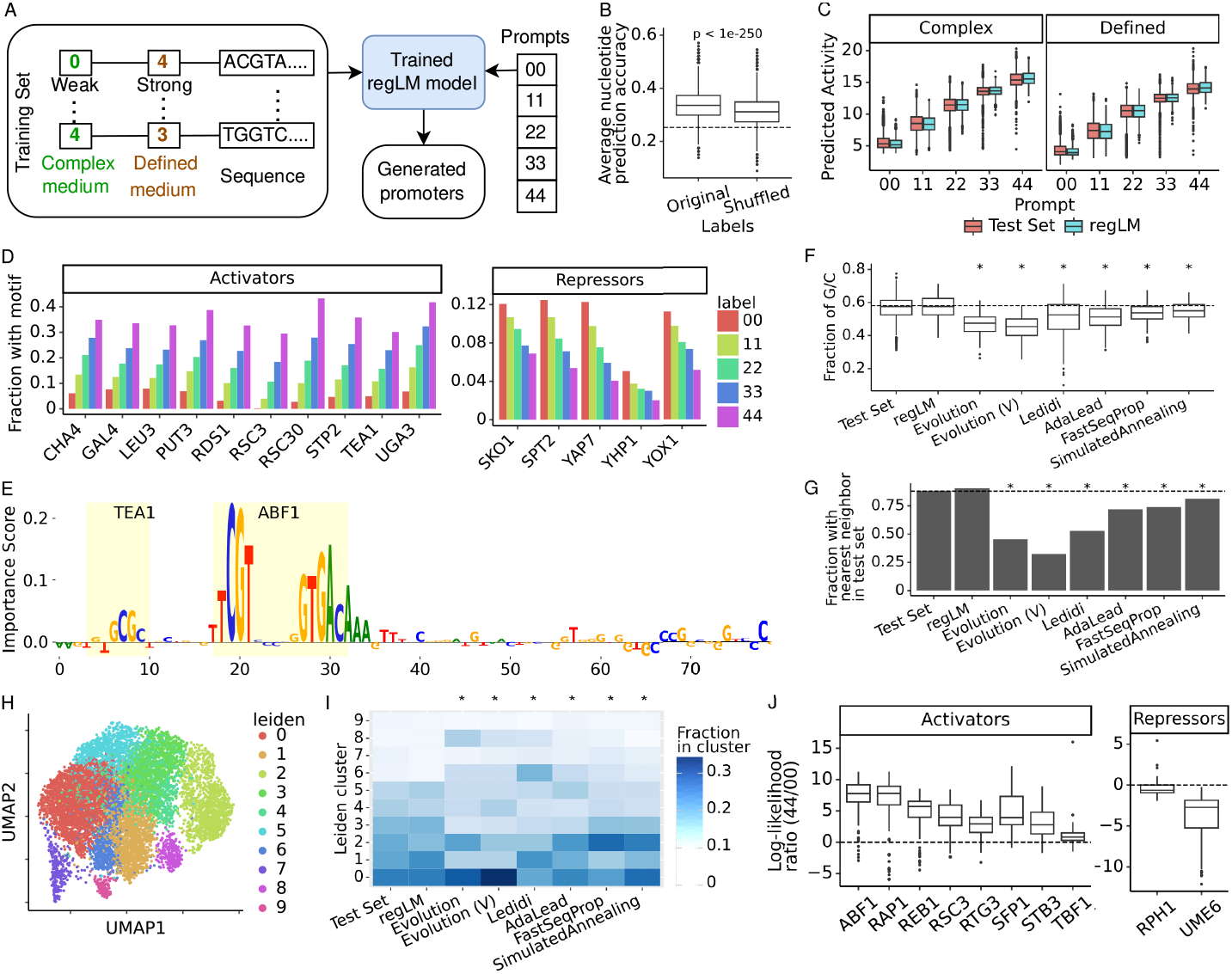
A) Schematic of the experiment. B) Boxplot showing the mean accuracy of the trained regLM model on test set sequences, before and after randomly shuffling the labels among sequences. The dashed line represents the accuracy of 0.25 expected by chance. C) Predicted activity of regLM generated promoters, compared to promoters from the test set with the same label. D) Fraction of regLM promoters prompted with different labels that contain the TF motifs most strongly associated with promoter activity in the test set. E) Example of a regLM generated strong promoter. Height represents the per-nucleotide importance score obtained from the paired regression model using ISM. Motifs with high importance are highlighted. F) Fraction of G/C bases in strong promoters generated by different methods. G) Fraction of generated promoters whose nearest neighbor based on k-mer content is a validated promoter from the test set, for different methods. H) UMAP visualization of real (Test Set) and synthetic strong promoters, labeled by cluster membership. I) Cluster distribution of strong promoters generated by different methods. J) Boxplots showing the log-ratio between the likelihood of the motif sequence given label 44 (high activity) vs. label 00 (low activity) for activating or repressing TF motifs inserted in random sequences. Motifs were selected based on TF-MoDISco results. In F, G, and I, asterisks indicate significant differences from the test set, and Evolution (V) represents synthetic promoters generated by Vaishnav et al. (2022) using Directed Evolution.

A regLM model trained to perform next nucleotide prediction on this dataset reached 31% mean accuracy on native yeast promoters and 33.8% accuracy on the test set, compared to 25% expected by chance (Fig. S2). Accuracy reduced when we randomly shuffled the labels across sequences (Fig. 2B, Fig. S3; One-sided Mann-Whitney U test p-value=8.8x10^−37^ for native promoters, p<10^−250^ for test set), indicating that the model learned to use the information encoded in the prompt tokens.

Within the test set, we observed higher accuracy in motifs for known yeast TFs (Fig. S4A; One-sided Mann-Whitney U test p-value=5.3x10^−77^). Accuracy increased with the abundance of the motif in the dataset (Fig. S4B, Pearson’s rho=0.54, p-value=1.2x10^−11^). Putting these observations together, we asked whether the model learned to associate specific motifs with categories of promoter activity. For each motif, we calculated the relative abundance of the motif in strong promoters (label 44) vs. weak promoters (label 00). We also calculated the ratio between the model’s accuracy within the motif when the motif was present in strong promoters vs. weak promoters. The strong correlation between these two metrics (Fig. S4C, Pearson’s rho=0.87, p-value=4.2x10^−42^), indicates that the model has learned to associate the prompt tokens with motifs that are consistent with the corresponding promoter activity; for example, when it observes the prompt 44, the model is more accurate at predicting motifs that tend to occur in strong promoters.

#### Generating synthetic yeast promoters

We generated promoters of defined strength by prompting the trained regLM model with labels 00, 11, 22, 33, and 44. Generated sequences were distinct from each other and from the training set, having a minimum edit distance of 25 bp from training sequences. Supervised regression models trained on the same data as the language model (Fig. S5) were used to discard generated sequences whose predicted activity did not match the prompt. Only 1.1% of the generated sequences were discarded.

Independent regression models trained on fully separate data from the language model (Fig. S6) predicted that regLM generates stronger promoters when prompted with higher labels, and that the activity of the generated promoters matches that of held-out test promoters with the same label (Fig. 2C). The abundance of TF motifs in the generated promoters was strongly correlated with their abundance in the test set; in other words, when regLM was prompted with the label 44 its generated sequences were more likely to contain motifs for activating yeast TFs that are often seen in strong promoters (Fig. S7, Fig. 2D).

In addition to motif abundance, we also examined per-base importance using *In silico* mutagenesis (ISM). Using TF-MoDISco [18], we identified motifs for known activator (ABF1, REB1, RAP1, RSC3, SFP1, STB3) and repressor (UME6) TFs with high importance in both the test set and the generated promoters, indicating that regLM generates motifs that contribute strongly to regulatory activity (Table S1, Fig. 2E; Fig. S8,9).

#### regLM generates promoters with diverse and realistic sequence content

To assess the biological realism of regLM-generated promoters relative to CREs generated by other methods, we compared 200 putative strong promoters generated by regLM (prompted with label 44) to sequences of similar predicted activity (Fig. S10) generated by five approaches (Directed evolution, Ledidi [17], AdaLead, FastSeqProp, and Simulated Annealing) as well as synthetic strong promoters generated in another study [21]. For a fair comparison, we performed all five model-guided methods using the regression model trained on the same dataset as regLM as an oracle. All sets of synthetic promoters were compared to known strong promoters from the test set using Polygraph [11]. Below, we use Evolution (V) to refer to synthetic promoters generated by Vaishnav et al. [21] using Directed Evolution.

GC content (the percentage of G or C nucleotides in a sequence) is a useful biological metric to evaluate the realism of synthetic sequences. regLM promoters were most similar to test set promoters in GC content, whereas other approaches produced sequences with lower GC content (Fig. 2F; Kruskal-Wallis p-value 5.7x10^−173^; Dunn’s post-hoc p-values 3.5x10^−69^ (Evolution vs. Test Set), 1.5x10^−70^ (Evolution (V) vs. Test Set), 2.3x10^−15^ (Ledidi vs. Test Set), 1.6x10^−27^ (AdaLead vs. Test Set) 4.2x10^−11^ (FastSeqProp vs. Test Set) 1.2x10^−6^ (Simulated Annealing vs. Test Set).

We counted the frequency of all k-mers of length 4 in all promoters. No k-mers were differentially abundant (defined as having two-sided Mann-Whitney U-test adjusted p-value < 0.05) in regLM promoters with respect to test set promoters, compared to 27-122 differentially abundant k-mers in the promoter sets generated by other methods (Table S2). When we matched each sequence to its nearest neighbor based on their k-mer frequencies, over 90% of regLM promoters were matched to a test set promoter, unlike other methods (Fig. 2G). regLM-generated promoters were among the most difficult to distinguish from the test set using simple classifiers based on k-mer frequency (Table S2).

We repeated the above analyses using the frequency of yeast TF binding motifs in all promoters (Methods). regLM and Simulated Annealing were the only methods that returned no differentially abundant motifs (defined as having two-sided Mann-Whitney U-test adjusted p-value < 0.05) with respect to the test set (Table S2). regLM-generated promoters were the most likely to have a test set promoter as their nearest neighbor based on motif frequency (Table S2). regLM-generated promoters were also among the most difficult to distinguish from the test set using simple classifiers based on motif frequency (Table S2).

To assess realism at the level of regulatory syntax, we examined combinations of motifs present in the generated sequences. We first computed the frequencies of pairwise combinations of motifs. Out of 2,321 motif pairs that were present in over 5% of any group of promoters, only 1 was differentially abundant (defined as having two-sided Fisher’s exact test adjusted p-value < 0.01) in regLM promoters with respect to test set promoters. In contrast, 21-439 motif pairs were differentially abundant in the other sets of synthetic promoters (Table S2). Motifs in regLM generated promoters also did not occur in significantly different positions compared to their positions in the test set (Table S2; defined as two sided Mann-Whitney U-test < 0.01).

We examined the distance and orientation between paired motifs in each group of promoters. For each motif pair, we counted the fraction of occurrences of the pair in which both motifs were in the same orientation, in each group of synthetic promoters as well as the test set. We found that the same-orientation fractions for motif pairs in regLM generated promoters showed the highest Pearson correlation with those in the test set (Table S2). We also tested whether the distance between motifs in these pairs was significantly different in synthetic promoters relative to the test set. regLM generated promoters were the third lowest in the number of motif pairs with significantly different distance (defined as having two-sided Mann-Whitney U-test adjusted p-value less than 0.01; Table S2).

To assess whether larger combinations of co-occurring motifs are shared between real and synthetic promoters, we performed graph-based clustering of real and synthetic strong promoters [20] based on their TF motif content. This revealed 10 clusters (Fig. 2H) corresponding to different combinations of co-occurring TF motifs (Fig. S11). All 10 clusters were represented in regLM promoters in similar proportion to their abundance in the test set; in contrast, the sets of sequences generated by other methods had skewed cluster representation, suggesting a tendency to converge upon specific transcriptional programs (Fig. 2I; Chi-squared p-values 2.6x10^−14^ (Evolution vs. Test Set), 3.6x10^−54^ (Evolution (V) vs. Test Set), 1.3x10^−68^ (Ledidi vs. Test Set), 1.0x10^−11^ (AdaLead vs. Test Set) 4.1x10^−3^ (FastSeqProp vs. Test Set) 2.8x10^−3^ (Simulated Annealing vs. Test Set).

Finally, we embedded all the real and synthetic promoters in a latent space defined by the convolutional layers of the independent regression models. The distance between sequences in this latent space incorporates not only differences in the frequency of important motifs, but also more complex regulatory syntax learned by the regression model such as motif orientation and spacing. Within this latent space, regLM promoters were still the most likely to have a test set promoter as their nearest neighbor (Table S2). Together, this evidence demonstrates comprehensively that regLM has learned many aspects of the yeast regulatory code.

#### Interrogating the trained regLM model reveals species-specific regulatory grammar

To learn whether interrogating the trained regLM model could reveal regulatory rules of yeast cells, we selected motifs for activating and repressing yeast TFs based on TF-MoDISco results (see Methods) and inserted each motif into 100 random DNA sequences. We used the trained regLM model to compute the likelihoods of the resulting sequences (P(sequence | label)) given either label 44 (strong promoter) or 00 (weak promoter). For each synthetic promoter, we defined a log-likelihood ratio as follows:

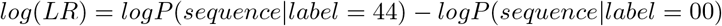

A positive log-ratio indicates that the model has learned the motif is more likely to occur in sequences with label 44 than 00, whereas a negative log-ratio indicates the opposite. We observed that sequences containing activating motifs tend to have positive log-likelihood ratios whereas sequences containing repressive motifs tend to have negative log-likelihood ratios (Fig. 2J).

We also calculated the per-base log-likelihood ratios on all promoters in the test set and found a significant positive correlation with the ISM scores derived from regression models (Fig. S12) further supporting our assertion that the language model has learned regulatory syntax, and suggesting that the log-likelihood ratio can be used as a nucleotide-level or region-level score to interpret these models.

### 2.3 regLM generates cell type-specific human enhancers

We trained a regLM model on a dataset of 200bp human enhancers and their measured activity in three cell lines (K562, HepG2 and SK-N-SH) [10] with the aim of designing cell type-specific human enhancers. Each sequence was prefixed with a sequence of 3 prompt tokens, each ranging from 0 to 3 and representing the measured activity of the enhancer in one of the 3 lines (Fig. S13). For example, label 031 indicates that the sequence has low activity in HepG2 cells, high activity in K562 cells, and weak activity in SK-N-SH cells (Fig. 3A).

**Fig. 3.**
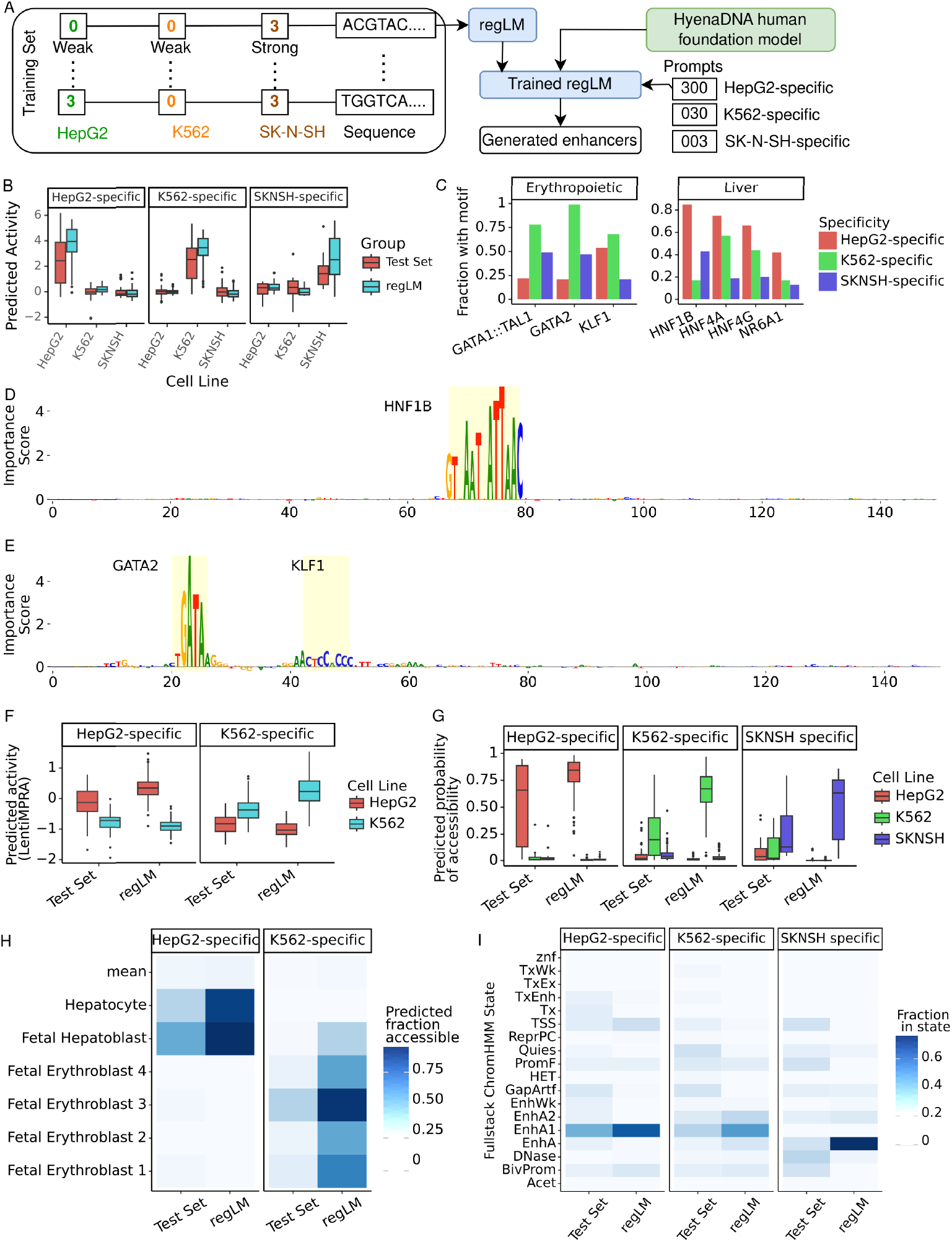
A) Schematic of the experiment. B) Predicted activity of cell type specific human enhancers generated by regLM, compared to real cell line-specific human enhancers from the test set, in 3 cell lines C) Fraction of regLM-generated enhancers containing selected cell type-specific TF motifs. D) Sequence of a HepG2-specific regLM-generated enhancer. E) Sequence of a K562-specific regLM-generated enhancer. Height is proportional to per-nucleotide importance scores from the independent regression model using ISM. Motifs with high importance are highlighted. F) Predicted activity of real and regLM-generated cell type-specific enhancers, using a model trained on LentiMPRA data. G) Predictions of a binary classification model trained on ATAC-seq from 3 cell lines, on real and regLM-generated cell type-specific enhancers. H) Predictions of a binary classification model trained on pseudobulk scATAC-seq from 203 cell types, on real and regLM-generated cell type-specific enhancers. Color intensity represents the fraction of sequences in the group that were predicted to be accessible. “mean” represents the average of all remaining cell types. I) Predictions of a classification model trained to classify genomic DNA into chromatin states defined by the fullstack chromHMM annotation, on real and regLM-generated cell type-specific enhancers. Color intensity represents the fraction of sequences in the group that were predicted to belong to the given state. Acet = acetylations, BivProm = bivalent promoter, EnhA, EnhA1, EnhA2 = Enhancers, EnhWk = Weak enhancers, GapArtf = Assembly gaps and artifacts, HET = heterochromatin, PromF = Flanking promoter, ReprPC= Polycomb repressed, Quies = Quiescent, TSS = Transcription start site, Tx = Transcription, TxWk = Weak transcription, TxEnh = Transcribed Enhancer, TxEx = Exon & Transcription, znf = ZNF genes

Here, instead of training a model from scratch, we could fine-tune a pre-existing HyenaDNA model that had already learned regulatory information from the human genome [15]. The trained model had a mean per-nucleotide accuracy of 45% on the test set. Despite the extreme rarity of cell type specific enhancers in the training set (enhancers with labels 300, 030 or 003 comprised only 0.16% of the training set), accuracy remained high on cell type specific enhancers (33.4%).

We used the trained regLM models to generate enhancers with activity specific to each cell line by prompting them with the labels 300 (HepG2-specific), 030 (K562-specific) and 003 (SK-N-SH specific). Generated sequences had a minimum edit distance of 21 nucleotides with reference to the training set. Using the regression models trained on the same dataset (Fig. S14) we selected the 100 regLM generated enhancers predicted to be most specific to each cell line.

Independent regression models (Fig. S15) predicted that the regLM generated elements are not only have cell type-specific activity, but in fact are more specific than the majority of enhancers with the corresponding label in the test set (Fig. 3B). In 2 of 3 cell types, regLM generated enhancers reach a level of specificity similar to previous synthetic cell type-specific enhancers [10] (Fig. S16), that were designed using model-guided approaches explicitly intended to maximize activity far beyond the range observed in the training set.

Since the test set contains very few enhancers with this level of cell type specificity, we did not evaluate the realism of the synthetic enhancers by quantitative comparisons of sequence content to the test set. However, we noted that motifs for known cell type-specific TFs occurred at higher frequency in regLM-generated enhancers of the appropriate specificity. For example, the motif for the erythropoietic TF GATA2 occurs at higher frequency in enhancers designed by regLM to be specific to K562 cells, whereas motifs for the liver-specific HNF4A and 4G factors occur at higher frequency in HepG2-specific synthetic enhancers (Fig. 3C). TF-MoDISco on per-base ISM scores yielded common motifs for cell type-specific TFs in the test set and in the generated enhancers (HNF1B, HNF4G, HNF4A for HepG2, and GATA2 for K562; Fig. 3D, E; Fig. S17, 18). We did not observe motifs for neuron-specific TFs among the TF-modisco outputs for SK-N-SH cells but instead general enhancer-associated factors such as AP-1.

We used several independent models to further validate the predicted cell type specificity of the HepG2- and K562-specific synthetic enhancers. First, we trained a regression model on lentiviral MPRA data from HepG2 and K562 cell lines [1], and applied it to our designed enhancers. The model predicted that the designed enhancers for K562 and HepG2 would still have cell line specific activity even in the context of lentiviral integration (Fig. 3F). Next, we trained binary classification models on chromatin accessibility data [6] from cell lines and predicted that the designed elements would have cell type specific chromatin accessibility (Fig. 3G). In addition, a model trained on chromatin accessibility in numerous fetal and adult human cell types predicted that the designed elements would also maintain cell type-specific accessibility in related cell types (Fig. 3H). Finally, we trained a model trained to classify DNA elements into chromatin states defined by the ChromHMM full-stack annotation [22]. This model predicted that most of the regLM generated enhancers belong to enhancer-associated chromatin states (Fig. 3I). In all cases, these models supported our prediction that that the cell type-specific enhancer activity of the regLM generated set exceeds that of cell type-specific enhancers in the test set.

We also ran all of these models on synthetic elements designed by [10] selected to have similar predicted activity to the regLM generated enhancers (Fig. S19), and found that the regLM generated enhancers showed comparable cell type specificity based on all predictions (Fig. S20, S21, S22, S23). Together, these diverse predictions greatly increase our confidence in the validity of regLM-generated enhancers.

## 3 Discussion

We demonstrate that the regLM framework successfully learns the regulatory code of DNA in different species and cell types, and generates diverse, realistic CREs with desired levels of activity *in silico*. Evaluation of synthetic sequences shows high concordance between the regulatory rules implemented in the sequences and known regulatory syntax. In the future, generated sequences can be experimentally validated to assess their function and safety.

While realistic sequences may help ensure predictable behavior in the genomic context, one weakness of our approach is that language models may learn correlated features in natural genomes that are not actually necessary for regulatory function. This can reduce functionality and be a weakness for mechanistic understanding. Larger training sets including randomly generated, mutated and non-genomic sequences will help mitigate this problem [3].

In conclusion, regLM can be easily adapted for numerous other biological tasks, such as codon optimization for mRNA. Our work suggests many directions for future research, such as improving prompt engineering and model interpretation for this novel architecture.

## 4 Methods

### 4.1 Training regLM models

HyenaDNA[15] is a decoder-only, sequence-to-sequence architecture consisting of a stack of blocks. Each block comprises a Hyena operator [16], followed by normalization and a feed-forward neural network. For human enhancers, we finetuned the pre-trained foundation model ‘hyenadna-medium-160k-seqlen’. This model has 6.55 million parameters and is trained to perform next token prediction on the human genome. For yeast promoters, we trained from scratch a Hye-naDNA model with the same architecture as ‘hyenadna-medium-160k-seqlen’.

For yeast promoters, the HyenaDNA model was trained for 100 epochs on 1 NVIDIA A100 GPU using the AdamW optimizer with cross-entropy loss, learning rate of 3x10^−4^ and batch size of 2048. Validation loss and accuracy were computed every 2000 steps and the model with highest validation accuracy was saved.

For human enhancers, the pre-trained HyenaDNA model was fine-tuned for 16 epochs on 1 NVIDIA A100 GPU using the AdamW optimizer with cross-entropy loss, learning rate of 10^−4^ and batch size of 1024. Validation loss and accuracy were computed every 100 steps and the model with highest validation accuracy was saved. During training, examples with each label were sampled from the training set with a weight inversely proportional to the frequency of the label, allowing the model to focus on cell type-specific enhancers that were extremely rare.

### 4.2 Generating synthetic CREs using regLM

#### Yeast promoters

We prompted the regLM model trained on yeast promoters to generate 1000 sequences each with labels 00, 11, 22, 33, and 44. During generation, we applied nucleus sampling [12] with a top-p cutoff of 0.85 to increase the reliability of generation. Generated promoters were filtered using the regression model trained on the same data as the language model. For each medium, we first used the regression model to predict the activity of all sequences in its training set, and computed the mean and standard deviation of predicted activity for training sequences with each class token (0, 1, 2, 3, and 4). We then used the same model to predict the activity of all generated promoters in both media. We discarded generated promoters whose predicted activity in either medium was more than 2 standard deviations from the mean predicted activity of promoters with the same token in the training set. We performed this procedure separately for complex and defined media. We then randomly selected 200 synthetic promoters of each generated class (00, 11, 22, 33, and 44) to compare with other methods.

#### Human Enhancers

The regLM model trained on human enhancers was prompted to generate 2500 sequences each with tokens 300 (HepG2-specific), 030 (K562-specific), and 003 (SK-N-SH specific) and nucleus sampling with top-p cutoff 0.99. Generated enhancers were filtered using a regression model trained on the same data. We first filtered the generated sequences using absolute thresholds consistent with the prompted labels (predicted activity greater than 4 in the target cell type and less than 0.2 in the off-target cell type). Next, we estimated the cell type specificity of each sequence as the difference between its predicted activity in the target cell type and its maximum predicted activity in off-target cell types. Based on this, we selected the 100 most specific regLM-generated enhancers for each cell type.

### 4.3 *In silico* evaluation of synthetic CREs

#### K-mer content

The frequency of all subsequences of length 4 (4-mers) was counted in each real or synthetic promoter. Each sequence was thus represented by a 256 dimensional vector. To calculate the fraction of real nearest neighbors, we matched each sequence to its nearest neighbor out of all real and synthetic sequences. For each group of synthetic CREs, we calculated the proportion of sequences whose nearest neighbor was an experimentally validated CRE from the test set. To compute classification performance, we trained a Support Vector Machine (SVM) with 5-fold cross-validation to distinguish each set of synthetic sequences from the reference set based on their k-mer frequencies. The Area Under the Receiver Operator Curve (AUROC) for each SVM was reported as a measure of classification performance.

#### Transcription factor motif content

Position Probability Matrices (PPMs) were downloaded from the JASPAR 2024 database in MEME format. 170 PPMs for yeast were selected using the filters Species=“Saccharomyces cerevisiae” and Versions=“Latest version”. 755 PPMs were selected for humans using the filters Species=“Homo sapiens” and Versions=“Latest version”.

Pairwise correlations between motifs were also downloaded from the JASPAR 2024 database. Motifs were clustered based on their pairwise Pearson correlations using agglomerative clustering with a distance threshold of 0.1. For clusters consisting of 2 motifs, the motif with higher information content was chosen as the cluster representative and the other was discarded. For clusters containing more than 2 motifs, the motif that had the highest average Pearson correlation to the other cluster members was selected as the representative and the others were discarded. This resulted in a filtered set of 140 motifs for yeast and 464 for human.

Reading the MEME files, conversion of PPMs to PWMs and sequence scanning were performed using the pymemesuite package, with a uniform background frequency, default pseudocount of 0.1, and p-value threshold of 0.001. Each sequence was represented by the 140-dimensional vector of its motif frequency for all motifs. Each sequence was matched to its nearest neighbors in this vector space and the proportion of real nearest neighbors for each group of synthetic CREs was calculated as described above. Classification performance was calculated as described above.

#### Model-based embeddings

Real and synthetic CREs were embedded in a model-defined latent space by passing them as input to the model and taking the output of the convolutional tower. For yeast promoters, the embeddings from each regression model had 384 features. We concatenated the embeddings from the models trained on two media, resulting in an embedding vector of size 768 for each sequence. Values were clipped to the 1st and 99th percentiles of the distribution to remove extreme values. In order to compute nearest neighbors efficiently, we reduced the number of features to 50 using Principal Component Analysis (PCA). Each sequence was matched to its nearest neighbors in PCA space and the proportion of real nearest neighbors for each group of synthetic CREs was calculated as described above.

### 4.4 Interpretation of the regLM model trained on yeast promoters

regLM is trained to perform next token prediction, i.e. for each position in a DNA sequence, regLM predicts the probability of all possible bases (A, C, G and T) conditioned on the previous bases as well as the initial label. Thus, we can obtain the likelihood of an observed sequence conditioned on its initial label (P(sequence | label)) as the product of probabilities of the base observed at each position.

To assess whether regLM has learned the function of a given motif, we generated 100 random DNA sequences and inserted the consensus sequence for the motif at the center of each. We prefixed each sequence with label 00 (low activity) and used the trained regLM model to predict the probability of each base in the motif. We calculated the likelihood of the motif conditioned on the sequence being labeled with 00 (*P* (*sequence* | *label* = 00)). We then prefixed all 100 sequences with the label 44 (high activity in both media) and repeated the procedure, calculating the likelihood of the motif conditioned on the sequence being labeled with 44 (*P* (*sequence*|*label* = 44)).

We refer readers to the Supplementary Methods for more details on data processing, model training, model parameters, and running benchmark methods.

## 5 Code Availability

regLM is available at https://github.com/Genentech/regLM along with documentation and a tutorial for use. Model weights and code to perform the experiments in this paper are linked to from the GitHub repository. Experiments were performed using Python v3.8, PyTorch v1.13.0 and PyTorch Lightning v1.8.2.

