## Supplementary Material for "regLM: Designing realistic regulatory DNA with autoregressive language models"

#### 1 Supplementary Methods

##### 1.1 Data Sources

Yeast promoter data from [5] was downloaded from https://zenodo.org/record/4436477. The data consisted of randomly generated DNA sequences of approximately 80 bp length and their measured promoter activity in yeast cells, based on reporter gene expression in a Gigantically Parallel Reporter Assay (GPRA). Measurements for 31,349,363 sequences were available in complex medium, and measurements for 21,037,407 sequences were available in defined medium.

Human Massively Parallel Reporter Assay (MPRA) data was downloaded from the supplementary material of [4]. This dataset contains 798,064 enhancer sequences from the human genome, of approximately 200 bp length, along with their measured enhancer activity in three human cell lines: HepG2 (a liver carcinoma cell line), K562 (an erythroleukemia cell line), and SK-N-SH (a neuroblastoma cell line).

##### 1.2 Data Processing

**Yeast promoters** We removed the constant sequences flanking each promoter sequence and selected promoter sequences that were 80 bp long and contained no N characters. This left measurements for 23,414,517 sequences in the complex medium and measurements for 16,799,784 sequences in the defined medium.

We then split the dataset into 7,533,156 sequences whose activity was measured in both media, and sequences whose activity was measured in only one medium (15,881,361 sequences measured only in complex medium and 9,266,628 sequences measured only in defined medium).

Taking the 7,533,156 sequences with measured activity in both media, we randomly split them into training (7,483,156 sequences), validation (50,000 sequences), and test (50,000 sequences) sets. We calculated the quintiles (five equally sized bins) of measured activity levels in complex medium and defined medium separately based on the training set. We assigned each sequence in the training set a token 0-4 based on its quintile of activity in the complex medium, and a second token 0-4 based on its quintile of activity in the defined medium. 0 indicates that the sequence belongs to the lowest quintile and 4 indicates that the sequence belongs to the highest quintile, which we describe as a strong promoter. Thus, each sequence was assigned a label consisting of two categorical tokens. For example, label 00 means a sequence that is in the 1st (lowest) quintile of activity in both media. 40 means a sequence that is in the 5th (highest) quintile of activity in complex medium but the 1st in defined medium.

Sequences in the validation and test sets were also assigned labels based on the quintiles calculated on the training set. Each DNA sequence was prefixed with its assigned label.

**Human enhancers** We assigned each sequence in the training set a token 0-3 based on its activity in HepG2 cells, a second token 0-3 based on its activity in K562 cells, and a third token 0-3 based on its activity in SK-N-SH cells. 0 indicates activity less than 0.2, 1 indicates activity between 0.2 and 0.75, 2 indicates activity between 0.75 and 2.5, and 3 indicates activity greater than 2.5. These cutoff values roughly correspond to the 25th, 75th and 95th percentiles of activity.

Thus, each sequence was assigned a label consisting of three categorical tokens. For example, label 301 means an enhancer that is in the highest group in HepG2 cells, the lowest group of activity in K562 cells, and the 2nd lowest group in SK-N-SH cells. Each DNA sequence was prefixed with its assigned label.

We split the dataset by chromosome. We held out 94,451 sequences from chromosomes 7, 13, 21 and 22 to train independent regression models. Of these, sequences from chromosome 21 were used for validation while the remaining sequences were used for training.

We used the remaining 669,233 sequences to train the regLM model and its paired regression models. We randomly sampled 50 sequences with cell type-specific activity to use as a validation set while the others were used for training. The training set for regLM was used as a test set for the independent regression models whereas the training set for the independent regression models was used as a test set for regLM and its paired regression models.

##### 1.3 Regression models

Regression models were trained to take as input a one-hot encoded CRE sequence and predict as output its activity. All regression models were based on the Enformer architecture [2] and were built using the enformer-pytorch package (https://github.com/lucidrains/enformer-pytorch). All regression models were trained for 10 epochs on 1 NVIDIA A100 GPU using the Adam optimizer. Performance was measured as the Pearson correlation between measured and predicted CRE activity on the test set.

For both yeast and human datasets, we trained two sets of regression models. One set of models was trained on the same sequences as the regLM model (without categorical labels), to use in conjunction with regLM to filter and prioritize the generated sequences. A second set of models was trained on independent data that was held out from regLM, kept entirely separate from all generative methods and was used only for independent *in silico* validation of synthetic CREs. Within each set, we trained separate regression models for each medium or cell type. Hence, for yeast, we trained a total of 4 regression models (two each for the two media) and for humans, we trained a total of 6 (2 each for 3 cell types).

**Yeast promoters** All regression models were Enformer-based models with 3 convolutional blocks followed by 1 transformer encoder layer. The first convolutional block has 384 channels. Each model has a single output head that predicts promoter activity in one of the two media. Models were trained with learning rate 5 x 10-4 and batch size 2048. Validation set loss was measured after each epoch and the model with lowest validation loss was saved.

The regLM-matched models were all trained using the same training, validation and test data used to train regLM. For the independent regression models, the 21,609,084 sequences whose activity was measured only in complex medium and 11,460,087 sequences whose activity was measured only in defined medium were used. In each medium, 50,000 randomly chosen sequences were held out for validation and 50,000 were held out for testing. The remaining sequences were used for training.

**Human enhancers** For human data, we downloaded the pre-trained Enformer model and reduced its size by dropping the last 8 transformer encoder layers (leaving 7 convolutional blocks and 3 transformer encoder layers). For each of our regression models, we added a single output head that predicts the total measured expression for the input sequence in a specific cell type.

For the regLM-matched regression models, we fine-tuned the model on the same sequences as regLM. These models were fine-tuned with learning rate 10-4, batch size 1024, and MSE loss. During training, examples with each label were sampled from the training set with a weight inversely proportional to the frequency of the label, allowing the model to focus on cell type-specific enhancers that were extremely rare. Validation set loss was measured after each epoch and the model with lowest validation loss was saved.

##### 1.4 Generating synthetic yeast promoters with benchmark methods

In order to benchmark regLM against existing commonly used approaches, we ran five other methods to generate synthetic yeast promoters: Directed Evolution, Ledidi, AdaLead, FastSeqProp and Simulated Annealing. These are all model-guided methods that iteratively make edits to a starting sequence to maximize a defined objective function using a trained predictive model (the ’oracle’).

We randomly chose sequences that had been measured to have low activity in all conditions (label 00) as the starting sequences. To ensure a fair comparison to regLM-generated sequences, the regression models trained on the same data as regLM were used as oracles. All approaches were each run multiple times with a different starting sequence each time, to generate diverse synthetic CREs. We used the CODA software package [4] to run AdaLead, FastSeqProp and Simulated Annealing.

For yeast promoters, we aimed to generate promoters with high activity in both media. The objective function for all methods was the mean predicted activity in the two media. All methods were run 200 times, each time with a different initial sequence, resulting in a diverse set of synthetic promoters from each method. Parameters were tuned to achieve synthetic sequences with similar predicted activity to those generated by regLM.

The following parameters were used:
Directed Evolution: 10 iterations
Ledidi: max_iter=1000, l=20, lr=3x10-3
AdaLead: model_queries_per_batch=75
FastSeqProp: n_steps=5, learning_rate=0.1
Simulated Annealing: n_steps=220, n_proposals=5

For each method, each regLM-generated strong promoter was matched to the method-generated sequences that were closest to it in predicted activity (measured by the mean squared error across both conditions), resulting in a matched set of 200 putative strong promoters designed by each method. Thus, since the various groups of synthetic elements have highly similar predicted activity, we can compare their sequence content to assess which approach gives rise to more biologically realistic sequences while reaching the same objective.

##### 1.5 Additional Models for Human CREs

**Lentiviral MPRA model** Training data were derived from [1], specifically the three cell line experiment detailed in supplemental tables 6 (200bp sequences) and 7 (log-transformed MPRA values). We took the pre-trained Enformer model, dropped all but the first transformer layer, and fine-tuned the model on this dataset using a batch size of 512, learning rate of 1e-4, and MSE loss for 10 epochs with the Adam optimizer. Fine-tuning was restricted to autosomes with chromosome 10 used for validation and chromosome 11 for testing, excluding peaks overlapping ENCODE blacklist regions.

**ATAC-seq model** We downloaded public ATAC-seq data in the form of processed BAM files from the ENCODE project for the following cell lines: K562

(ENCFF534DCE), HepG2 (ENCFF624SON), GM12878 (ENCSR095QNB), IMR90 (ENCFF715NAV), WTC11 (ENCFF240QKT), and SK-N-SH (ENCFF270AGJ). As standard, SK-N-SH cells were treated with trans-retinoic acid prior to sequencing to induce neural-like differentiation. In addition, we downloaded raw fastq files for Jurkat cells from the short-read archive (SRX7785407). These raw reads were then aligned to hg38 following the ENCODEv4 standards.

Peaks for each cell line were called using MACS3 [8] callpeaks with the additional parameters "–nomodel –shift -100 –extsize 200". We then created a unified peak set as described in [3] with an SPM = 2 and an extension of 250bp. This resulted in a uniform peak set consisting of 366,776 500 bp regions to which we added an additional 15 percent of 500 bp regions overlapping no known peak to serve as no-signal background regions during modeling. These peak regions were binarized per cell line by overlapping them with cell line specific MACS3 peak calls using an SPM value of 5. Peaks were then resized to 200bp.

We took the pre-trained Enformer model, dropped all but the first transformer layer, and added a head (linear layer) with a sigmoid activation function to predict the probability of the input sequence being a peak in each cell line. We fine-tuned the model on this dataset using a batch size of 512, learning rate of 1e-4, and binary cross-entropy loss for 10 epochs with the Adam optimizer. Fine-tuning was restricted to autosomes with chromosome 10 used for validation and chromosome 11 for testing, excluding peaks overlapping ENCODE blacklist regions. Validation set loss was measured after each epoch and the model with lowest validation loss was saved. At inference, predicted probabilities were thresholded with a cutoff of 0.5 to generate final predictions.

**CATLAS scATAC-seq model** We downloaded binarized single-cell pseudobulk chromatin accessibility matrices from the CATLAS project [7]. Peaks were resized to 200 bp. Cell types with less than 3% positive labels were discarded which reduced the number of cell types from 222 to 203.

We took the pre-trained Enformer model, dropped all but the first transformer layer, and added a head (linear layer) with a sigmoid activation function to predict the probability of the input sequence being a peak in each cell type. We fine-tuned the model on this dataset using a batch size of 512, learning rate of 1e-4, and binary cross-entropy loss for 10 epochs with the Adam optimizer. Fine-tuning was restricted to autosomes with chromosome 7 used for validation and chromosome 13 for testing, excluding peaks overlapping ENCODE blacklist regions. Validation set loss was measured after each epoch and the model with lowest validation loss was saved. At inference, predicted probabilities were thresholded with a cutoff of 0.5 to generate final predictions.

**Full-stack chromHMM model** Annotations for the hg38 genome generated using the full-stack chromHMM model were obtained from [6]. These annotations were extended to 1024bp and restricted to the autosomes. The fine scale categories were collapsed by stripping off the prefix and suffix values to generate 18 broad categories of annotations (Acet (acetylations), BivProm (bivalent promoter), DNase, EnhA (Enhancers), EnhA1 (Enhancers), EnhA2 (Enhancers), EnhWk (Weak enhancers), GapArtf (Assembly gaps and artifacts), HET (heterochromatin), PromF (Flanking promoter), ReprPC (Polycomb repressed), Quies (Quiescent), TSS (Transcription start site), Tx (Transcription), TxWk (Weak transcription), TxEnh (Transcribed Enhancer), TxEx (Exon & Transcription), and znf (ZNF genes)). The resulting element set was downsampled to have a maximum of 200000 instances of any given category.

We took the pre-trained Enformer model, dropped all but the first transformer layer, and added a head with a Softmax activation function to predict class probabilities. We fine-tuned the model to perform multiclass classification on this dataset. For fine-tuning, we used a batch size of 512, learning rate of 1e-4, reverse complement augmentation, and cross-entropy loss for 14 epochs with the Adam optimizer. Fine-tuning was restricted to autosomes with chromosome 7 used for validation and chromosome 13 for testing, excluding regions overlapping ENCODE blacklist regions. Validation set loss was measured after each epoch and the model with lowest validation loss was saved. At inference, each sequence was predicted to belong to the class with the highest predicted probability.

#### 2 Supplementary Notes

##### 2.1 Choice of labels

In the above experiments, we generated labels by dividing yeast promoters into 5 equal bins per medium and dividing human enhancers into 4 unequal bins per cell type. In theory, any number of class labels can be used, and our package allows users to choose the number of labels and to define them in any way. However, the more we subdivide our data, the less information the model will have to learn an accurate distribution of each class. In contrast, having fewer subdivisions may make it more difficult for the model to share information across similar categories.

The resolution of the data should also be kept in mind. In case of the yeast promoter GPRA assay, promoters were sorted into bins based on their measured activity; however, many promoters were measured only once and so their measured values are not precise. Hence, we chose to use a lower resolution.

To demonstrate that our method can work with different labeling schemes, we have also re-trained the yeast model using 10 label classes instead of 5. The

10-class model performs well at generating sequences with label-consistent expression (Fig. S24).

In the case of human enhancers, the cell type specific enhancers we aimed to design were extremely rare in the dataset. Therefore instead of dividing the dataset into equally sized bins, we assigned sequences with extremely high activity (in the 95th percentile for each cell line) to a separate bin.

##### 2.2 Regulatory programs used by different methods

We clustered yeast promoters into groups containing different combinations of motifs and partitioned the synthetic promoters generated by each method into clusters. 23% of the strong promoters in the test set mostly fell into cluster 0, which is characterized by ABF1, SKO1, YAP6, and CIN5 motifs, followed by 18% in cluster 1 (RSC3, RSC30, DAL82, SUT1, and TEA1 motifs), 17% in cluster 2 (ECM22, HAL9, ERT1, CAT8, and ASG1 motifs) and 15% in cluster 3 (CHA4, PDR3, PDR1, IME1, and RDS1 motifs). regLM promoters partitioned similarly to test set promoters, with no significant differences. However, the most notable feature in all other methods was a strong enrichment for clusters 6 (CUP2, EDS1, STB3, SUM1, and SFP1 motifs) and 8 (ARR1, CIN5, FKH1, RLM1, and SPT15 motifs). In addition, promoters generated by FastSeqProp were enriched in clusters 2 (ECM22, HAL9, ERT1, CAT8, and ASG1 motifs) and 7 (characterized by the presence of XBP1 motifs).

### 3 Supplementary Tables

| | Transcription Factor | Motif | Contribution | TOMTOM q-value (test set) | TOMTOM q-value (regLM generated promoters) |
|---|---|---|---|---|---|
| 1 | ABF1 | MA0265.3 | Positive | $4.5 \times 10^{-8}$ | $4.4 \times 10^{-3}$ |
| 2 | REB1 | MA0363.3 | Positive | $1.2 \times 10^{-5}$ | $5.0 \times 10^{-4}$ |
| 3 | RAP1 | MA0359.3 | Positive | $1.9 \times 10^{-4}$ | $3.5 \times 10^{-5}$ |
| 4 | TBF1 | MA0403.3 | Positive | $3.4 \times 10^{-3}$ | N/A |
| 5 | RTG3 | MA0376.2 | Positive | $6.3 \times 10^{-3}$ | N/A |
| 6 | RSC3 | MA0374.2 | Positive | 0.03 | 0.02 |
| 7 | SFP1 | MA0378.2 | Positive | 0.04 | 0.01 |
| 8 | STB3 | MA0390.2 | Positive | 0.05 | 0.02 |
| 9 | UME6 | MA0412.3 | Negative | $1.4 \times 10^{-6}$ | $2.9 \times 10^{-4}$ |
| 10 | RPH1 | MA0372.2 | Negative | 0.014 | N/A |

**Table S1.** Motifs identified by TF-MoDISco and TOMTOM on test set and regLM generated promoters. N/A indicates that no motif with a significant match was found by TF-MoDISco.

| | Test Set | regLM | Evolution | Evolution (V) | Ledidi | AdaLead | FastSeq-Prop | Simulated Anneal-ing |
|---|---|---|---|---|---|---|---|---|
| Differential k-mers w.r.t. Test Set | N/A | **0** | 86 | 122 | 51 | 33 | 27 | 71 |
| Fraction of Nearest Neighbors in Test Set (k-mer frequency) | 0.88 | **0.91** | 0.45 | 0.33 | 0.53 | 0.73 | 0.73 | 0.87 |
| SVM AUROC vs. Test Set (k-mer frequency) | N/A | **0.5** | 0.52 | 0.64 | 0.57 | **0.5** | **0.5** | 0.51 |
| Differential motifs w.r.t. Test Set | N/A | **0** | 22 | 42 | 5 | 6 | 1 | **0** |
| Fraction of Nearest Neighbors in Test Set (motif frequency) | 0.85 | **0.83** | 0.67 | 0.47 | 0.71 | 0.75 | 0.82 | 0.82 |
| SVM AUROC vs. Test Set (motif frequency) | N/A | **0.5** | 0.52 | 0.53 | 0.53 | **0.5** | **0.5** | **0.5** |
| Differential motif pairs w.r.t. Test Set | N/A | **1** | 285 | 439 | 153 | 125 | 89 | 21 |
| Differential motif positioning w.r.t. Test Set | N/A | **0** | 1 | 9 | 2 | 3 | **0** | 2 |
| Pearson Rho (fraction of pair in same orientation) with test set | N/A | **0.52** | 0.51 | 0.34 | 0.47 | 0.45 | 0.45 | 0.41 |
| Differential inter-motif distance w.r.t. Test Set | N/A | 1897 | 1853 | **1846** | 2052 | 1909 | 1945 | 1968 |
| Fraction of Nearest Neighbors in Test Set (model embedding) | 0.88 | **0.88** | 0.80 | 0.53 | 0.79 | 0.81 | 0.86 | 0.86 |

**Table S2.** Additional metrics for all sets of synthetic yeast promoters as well as strong promoters from the test set. The best performances for each method are highlighted in bold. Evolution (V) represents synthetic promoters generated by [5] using Directed Evolution.

#### 4 Supplementary Figures

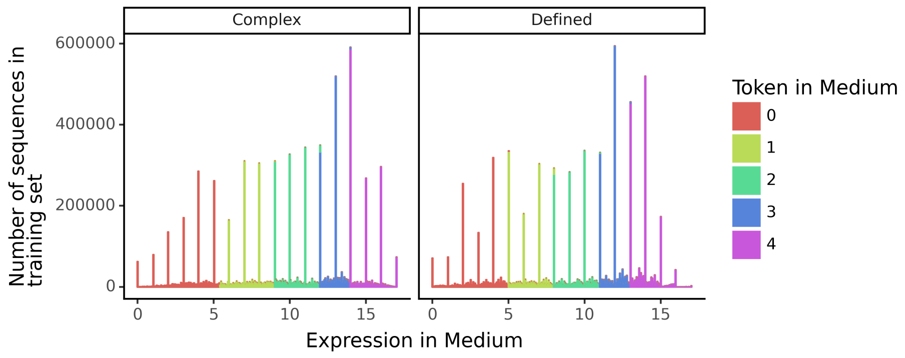


**Fig. S1.** Histograms of measured promoter activity in the two media, colored by the assigned token in each medium. Each promoter was assigned a token ranging from 0-4 where 0 corresponds to the lowest quintile of measured activity and 4 corresponds to the highest. This procedure was performed separately for measurements in complex and defined media.

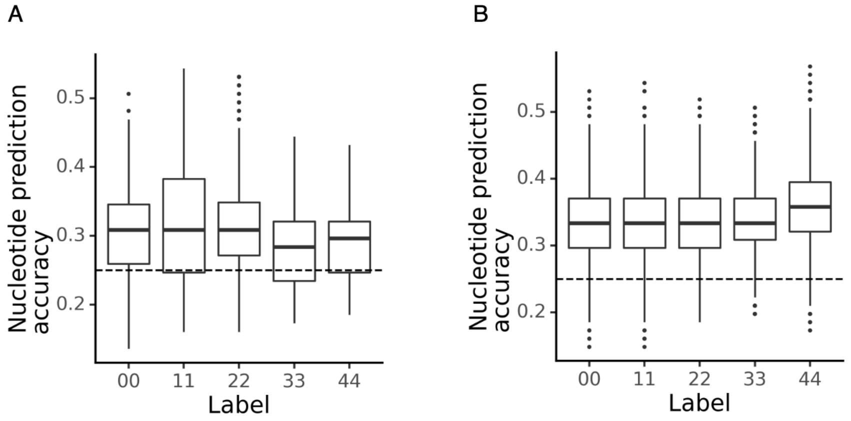


**Fig. S2.** A. Boxplots showing the average per-nucleotide prediction accuracy of the yeast regLM model on A) 3,922 native yeast promoters and B) 50,000 promoters from the test set, separated by the promoter class labels. Only the 5 most common labels (00, 11, 22, 33, 44) are shown. The dashed lines represent the accuracy of 0.25 expected by chance.

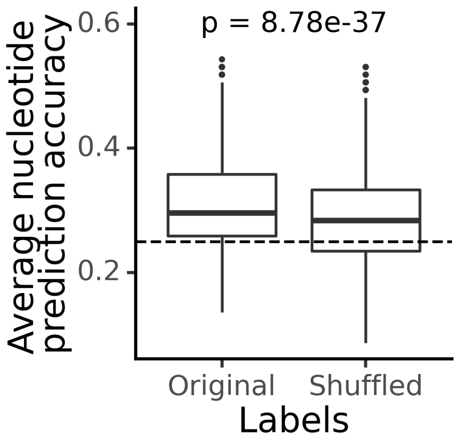


**Fig. S3.** Boxplots showing the average per-nucleotide prediction accuracy of the yeast regLM model on 3,922 native yeast promoters, before and after shuffling the labels across sequences. The dashed line represents the accuracy of 0.25 expected by chance.

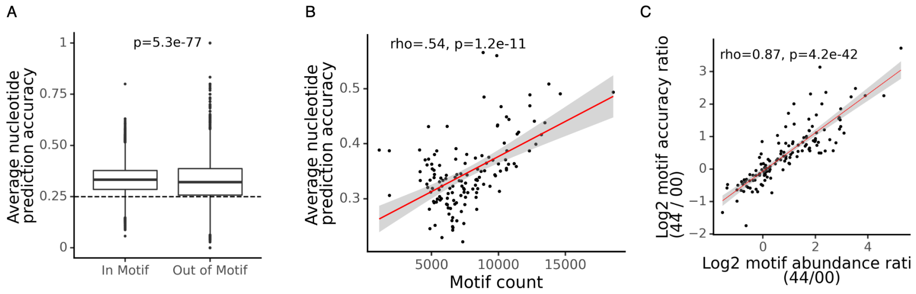


**Fig. S4.** A) Average nucleotide prediction accuracy of the regLM model on 50,000 promoters in the test set, for nucleotides within known TF-binding motifs versus those outside. The dashed line represents the accuracy of 0.25 expected by chance. B) Scatterplot showing the accuracy of the regLM model on nucleotides within TF binding motifs in the test set. Each point represents a TF binding motif. The x-axis shows the number of occurrences of the motif across the test set. The y-axis shows the average accuracy of the regLM model on all instances of the motif in the test set. C) Scatterplot showing the log ratio between the abundance of a motif in strong promoters (label 44) vs. weak promoters (label 00) on the x-axis, and the log ratio between the average accuracy of the regLM model on all instances of the motif in strong promoters vs. weak promoters on the y-axis. Red lines in B) and C) show the linear fit to the data.

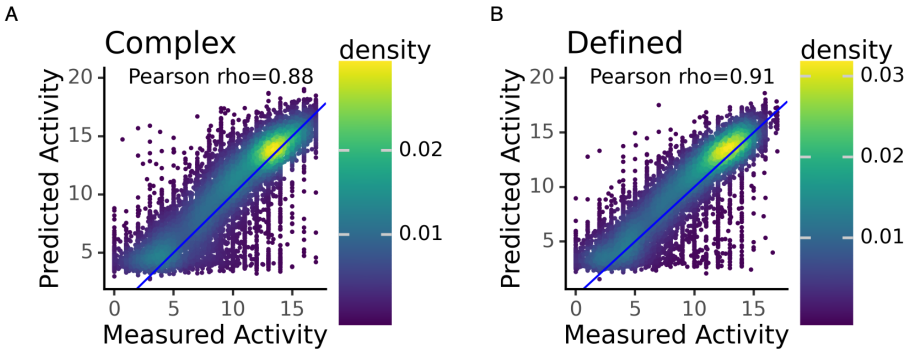


**Fig. S5.** Performance of supervised regression models trained to predict promoter activity of yeast promoter sequences, in A) complex medium and B) defined medium. The models were trained and tested on the same data as the regLM model. Scatterplots show the measured and predicted activity of 50,000 test set promoters.

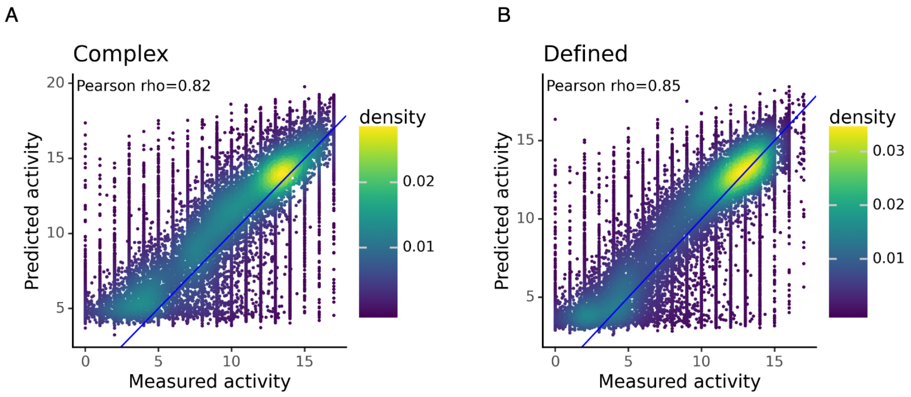


**Fig. S6.** Performance of two supervised regression models trained to predict promoter activity of yeast promoter sequences in A) complex and B) defined medium respectively. These models were trained and tested on separate data from the regLM model. Scatterplots show the measured and predicted activity of 50,000 test set promoters each.

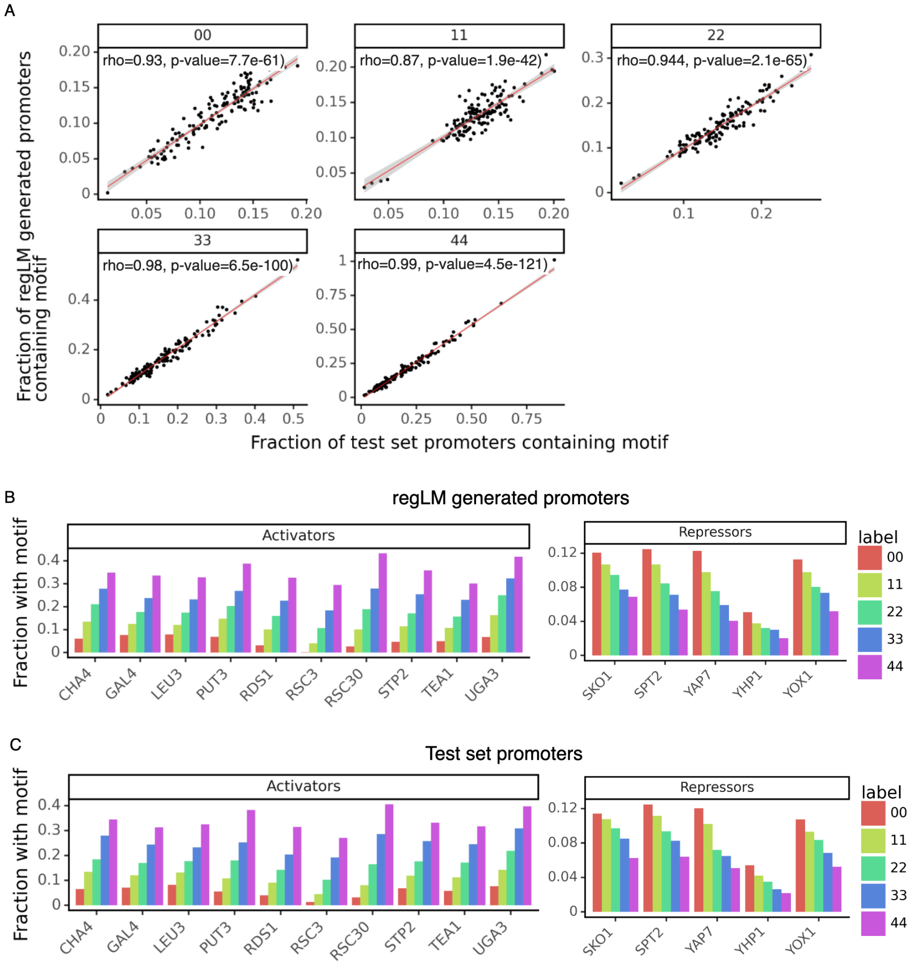


**Fig. S7.** A) Scatterplot showing the association between motif abundance in regLM generated promoters versus the test set, for promoters with different labels. Red lines show the linear fit to the data. B, C) A closer focus on the motifs that show the strongest differential abundance between strong and weak promoters in the test set, showing the close match between their abundance in the test set and in the generated promoters. Bar plots show the fraction of regLM generated promoters and test set promoters that contain selected activating and repressing TF motifs, separated by label. Only the 5 most common labels (00, 11, 22, 33, 44) are shown.

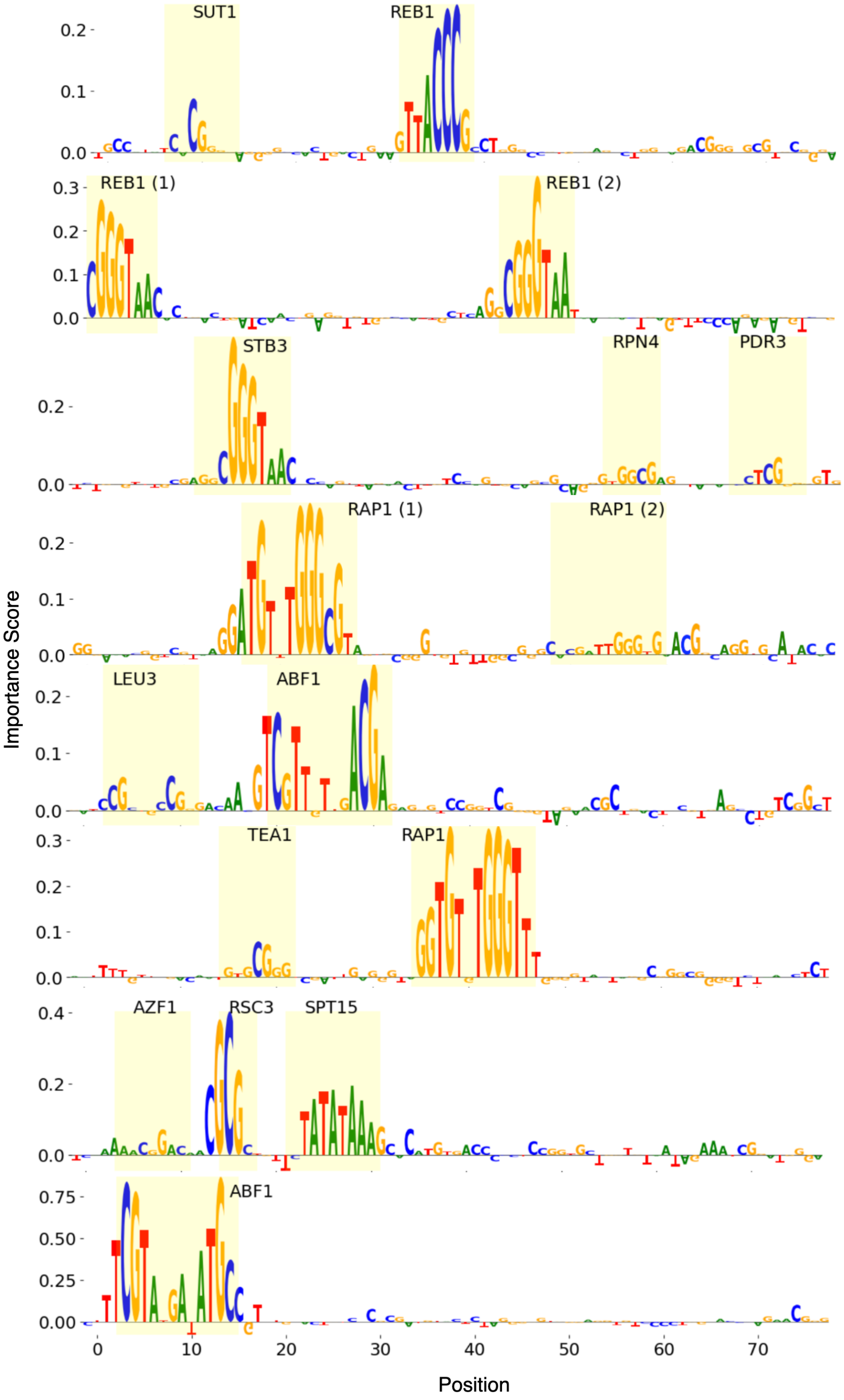


**Fig. S8.** Examples of strong promoters in the test set. Height represents the per-nucleotide importance score obtained from the paired regression model using ISM. Motifs with high importance are highlighted.

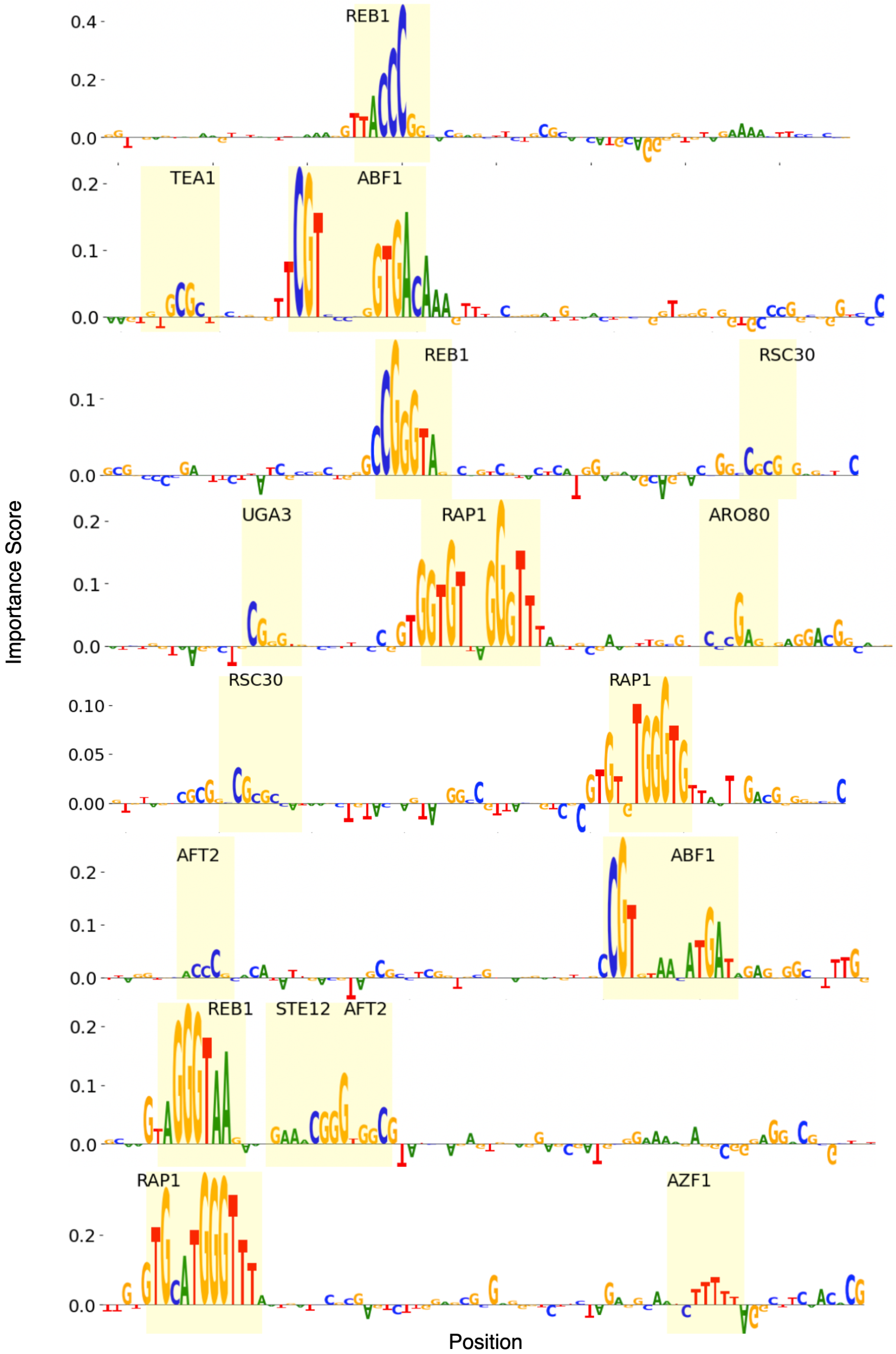


**Fig. S9.** Examples of strong promoters generated by regLM. Height represents the per-nucleotide importance score obtained from the paired regression model using ISM. Motifs with high importance are highlighted.

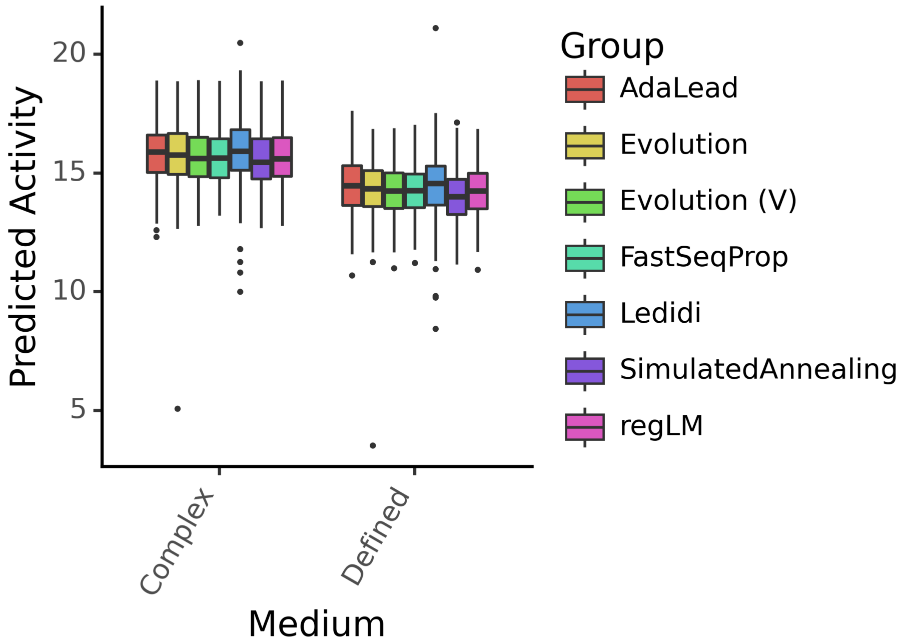


**Fig. S10.** Predicted activity of synthetic strong yeast promoters generated by different methods, in complex and defined media. 200 synthetic promoters were generated by each method. Evolution (V) represents synthetic promoters generated by Vaishnav et al. [5] using Directed Evolution.

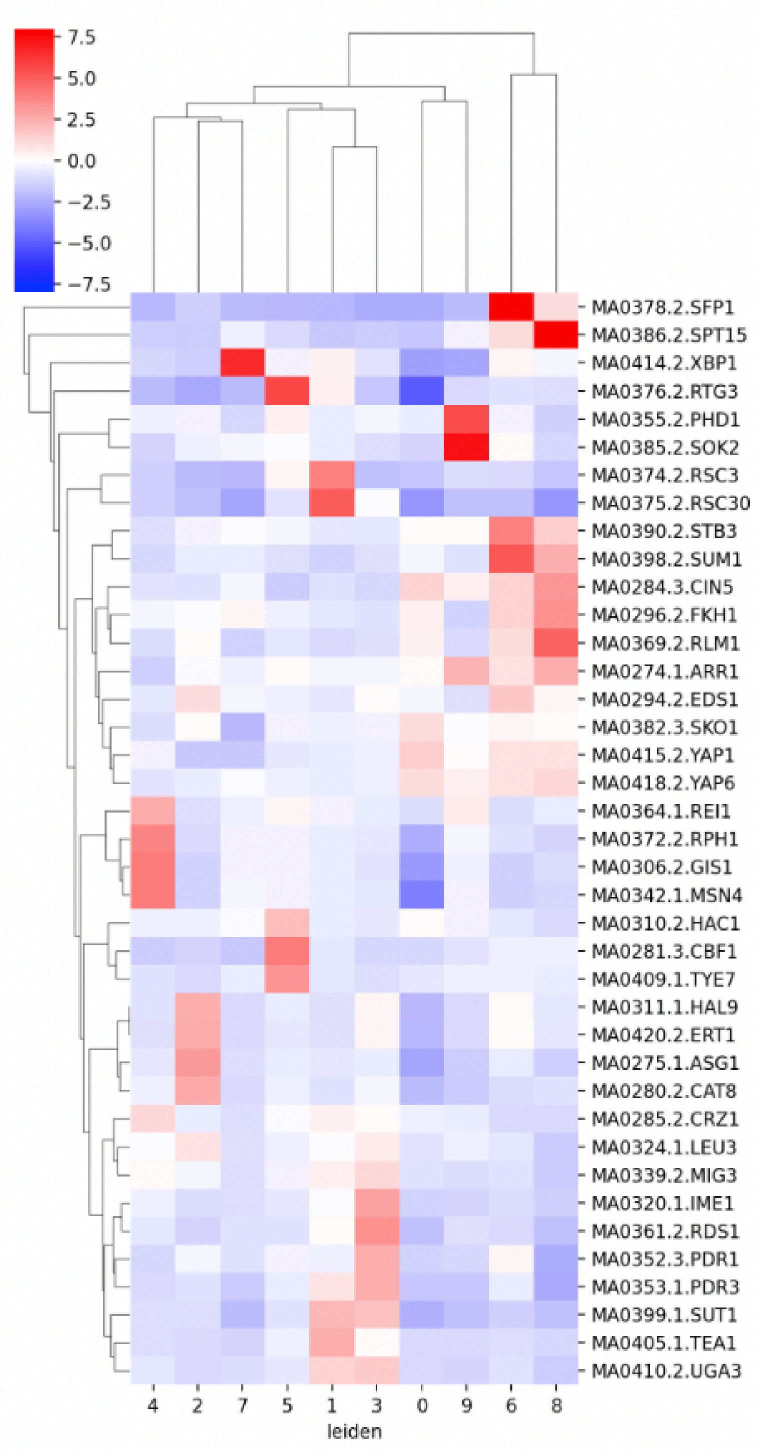


**Fig. S11.** Heatmap showing log2 fold changes in motif abundance for the top differentially abundant motifs across clusters of strong promoters. Fold changes were calculated for each cluster relative to the entire dataset.

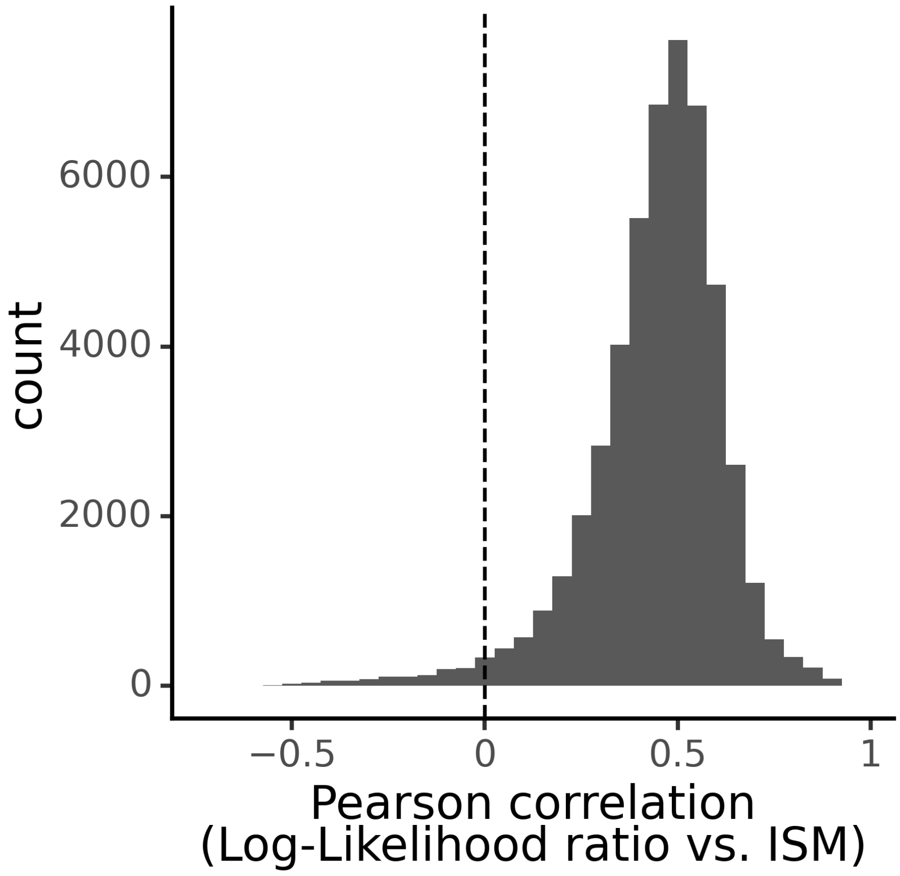


**Fig. S12.** Histogram of the Pearson correlation between per-base ISM scores from the paired regression model, and the per-base log-likelihood ratios from the regLM model (44 vs. 00), across all 50,000 promoters in the test set.

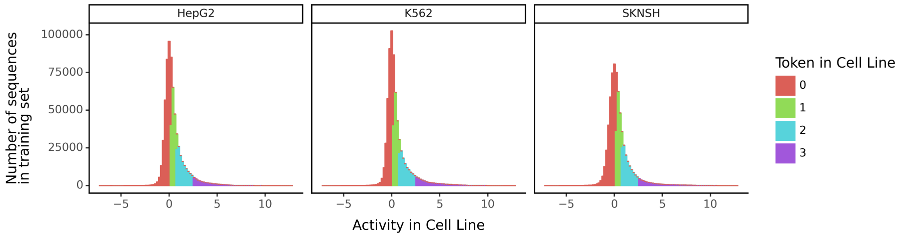


**Fig. S13.** Histograms showing the measured enhancer activity in each of 3 cell lines, for cell type-specific enhancers in the training set. Sequences were divided into 4 bins based on their measured activity. Each sequence was assigned a token ranging from 0-3 where 0 corresponds to the lowest bin and 3 corresponds to the highest. This procedure was performed separately for measurements in each cell line. The color corresponds to the assigned token in that cell line.

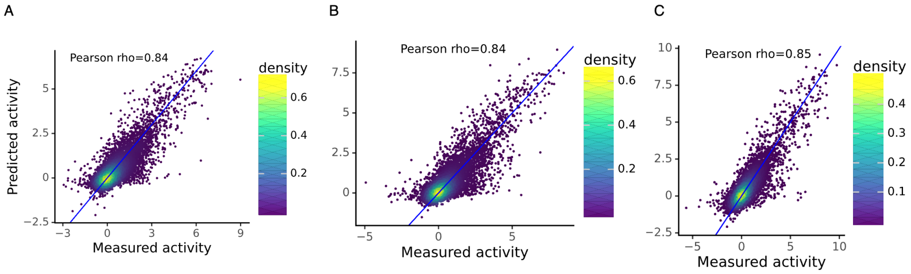


**Fig. S14.** Performance of a supervised regression model trained to predict activity of human enhancer sequences in A) HepG2 cells B) K562 cells C) SK-N-SH cells. The models were trained and tested on the same data as the regLM model. Scatter plots show the measured and predicted activity of enhancers in the test set.

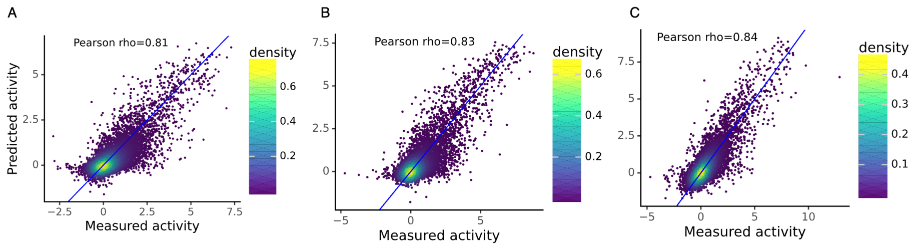


**Fig. S15.** Performance of a supervised regression model trained to predict activity of human enhancer sequences in A) HepG2 cells B) K562 cells C) SK-N-SH cells. The models were trained and tested on separate data from the regLM model. Scatter plots show the measured and predicted activity of enhancers in the test set.

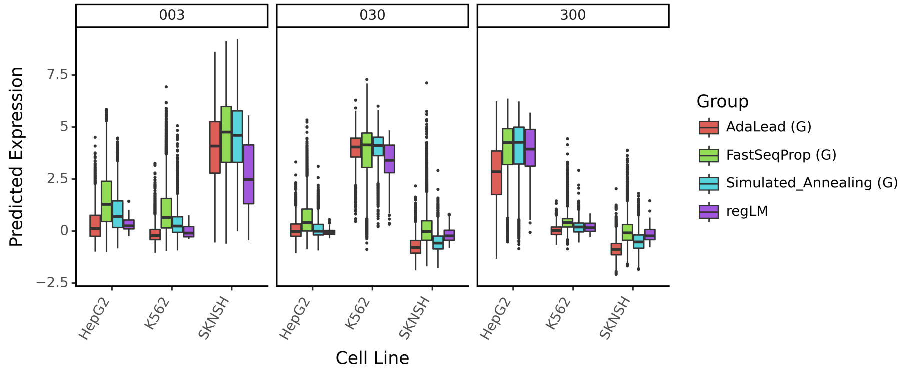


**Fig. S16.** Predicted activity of synthetic cell type-specific enhancers generated by regLM and by Gosai et al. [4], in 3 cell lines. Predictions were generated by regression models trained on separate data from regLM. (G) indicates that the method was performed by Gosai et al. [4]

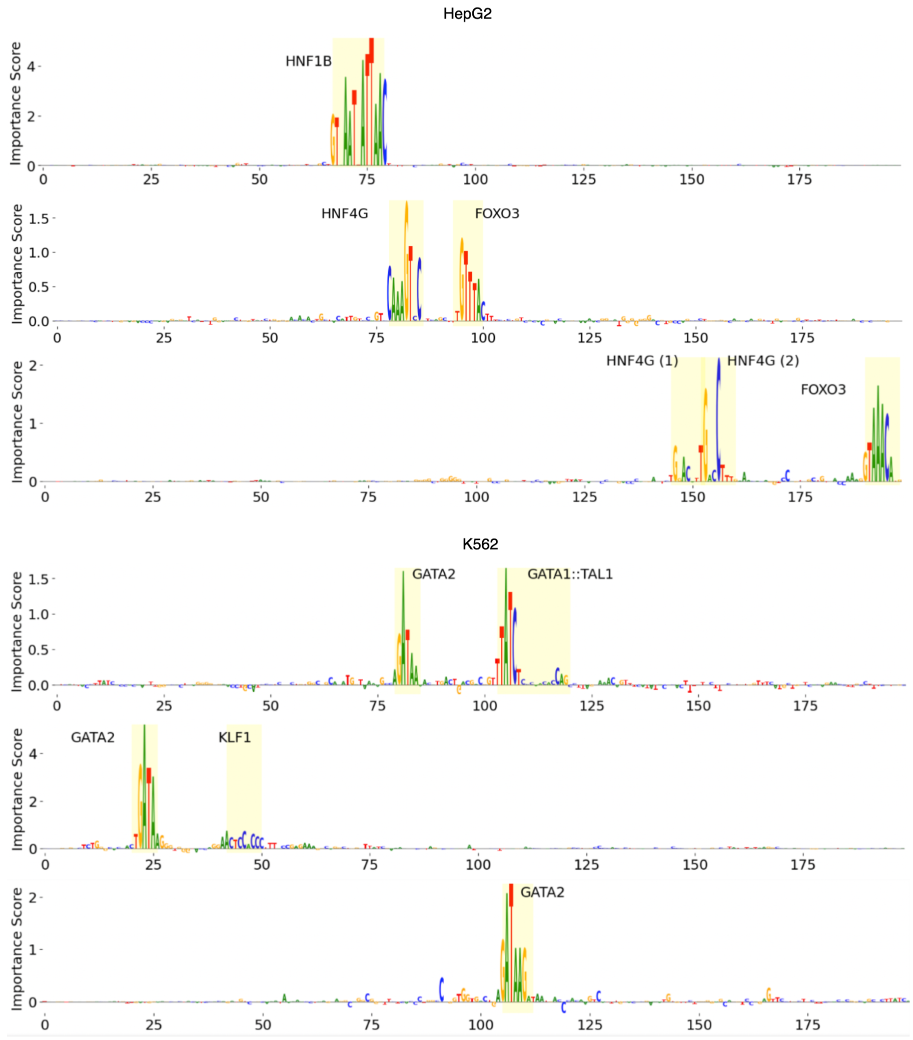


**Fig. S17.** ISM-based importance scores for regLM generated K562 and HepG2-specific enhancers, highlighting highly contributing motifs.

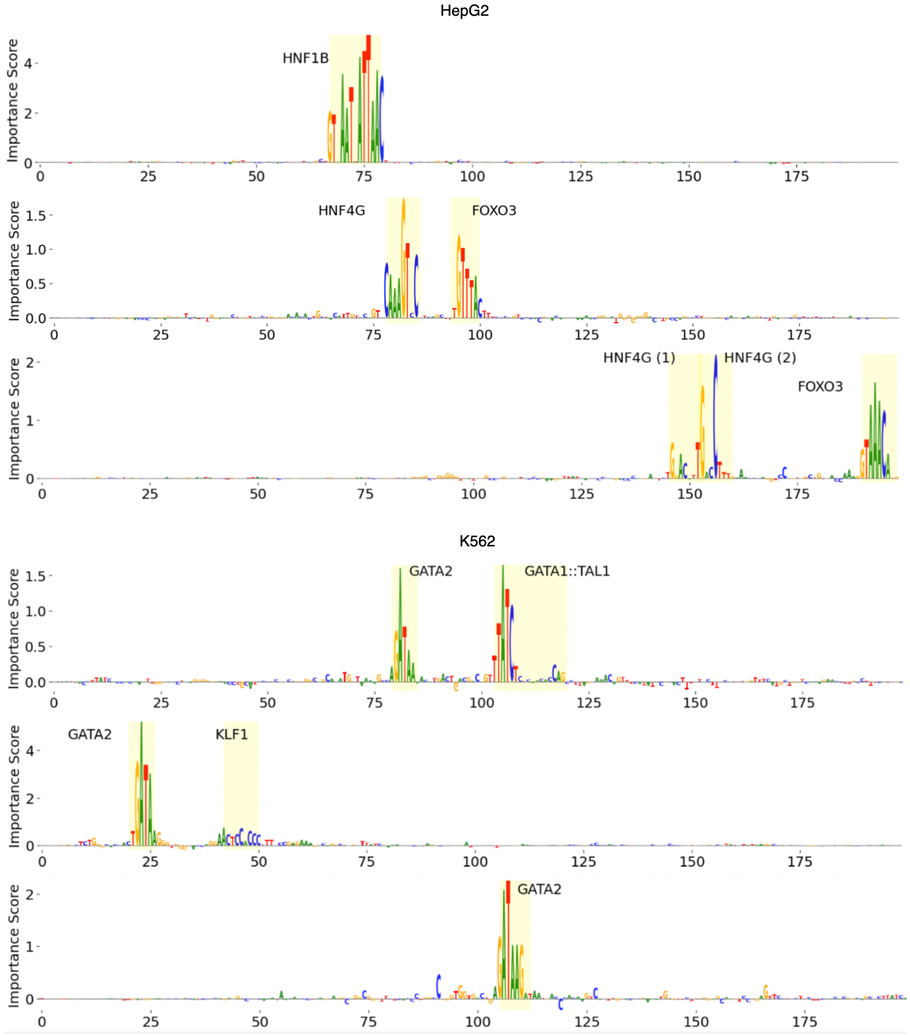


**Fig. S18.** ISM-based importance scores for K562 and HepG2-specific enhancers in the test set, highlighting highly contributing motifs.

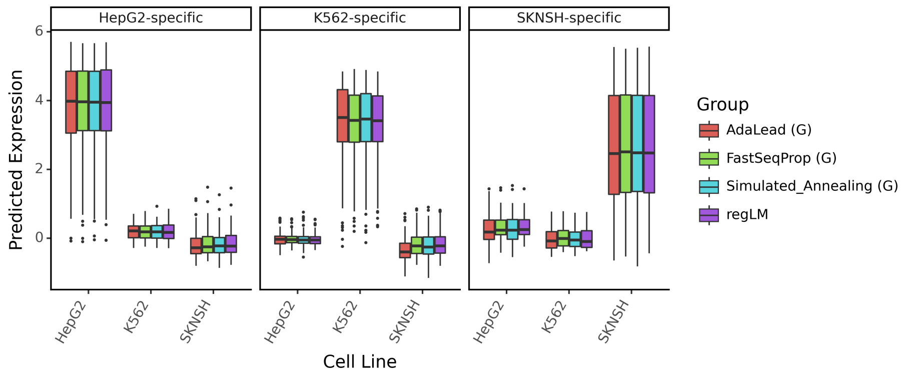


**Fig. S19.** Predicted activity of 100 synthetic cell type-specific enhancers generated by regLM for each cell line, and the 100 Gosai et al. [4] designed elements chosen to have the most similar activity for each cell line. (G) indicates that the method was performed by Gosai et al. [4]

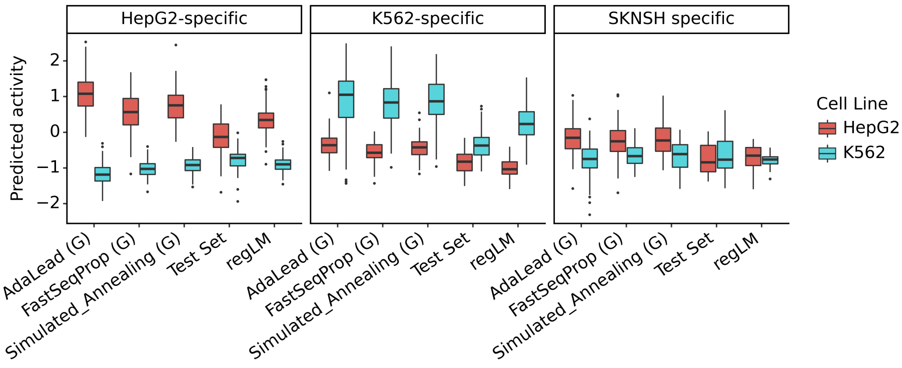


**Fig. S20.** Predicted activity of synthetic cell type-specific enhancers generated by different methods, using a model trained on Lentiviral MPRA data. (G) indicates that the method was performed by Gosai et al.[4]. The 300 Gosai et al. [4] designed elements chosen based on similar activity to regLM generated enhancers are shown here.

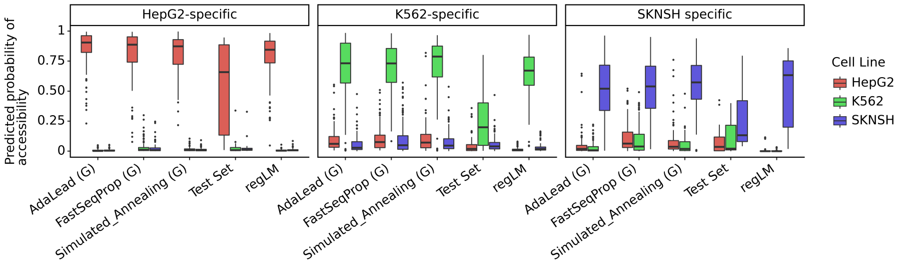


**Fig. S21.** Predictions of a binary classification model trained to predict ATAC-seq peaks in three cell lines, on synthetic cell type-specific enhancers generated by different methods. (G) indicates that the method was performed by Gosai et al.[4]. The 300 Gosai et al. [4] designed elements chosen based on similar activity to regLM generated enhancers are shown here.

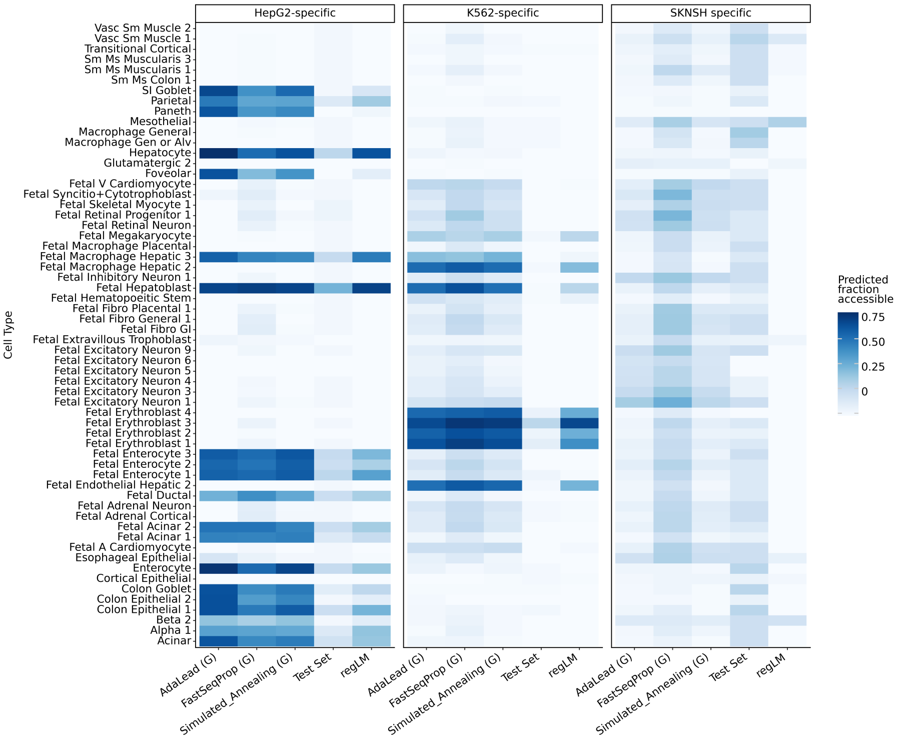


**Fig. S22.** Predictions of a binary classification model trained to predict ATAC-seq peaks in 203 cell types, on synthetic cell type-specific enhancers generated by different methods. (G) indicates that the method was performed by Gosai et al.[4]. The 300 Gosai et al. [4] designed elements chosen based on similar activity to regLM generated enhancers are shown here.

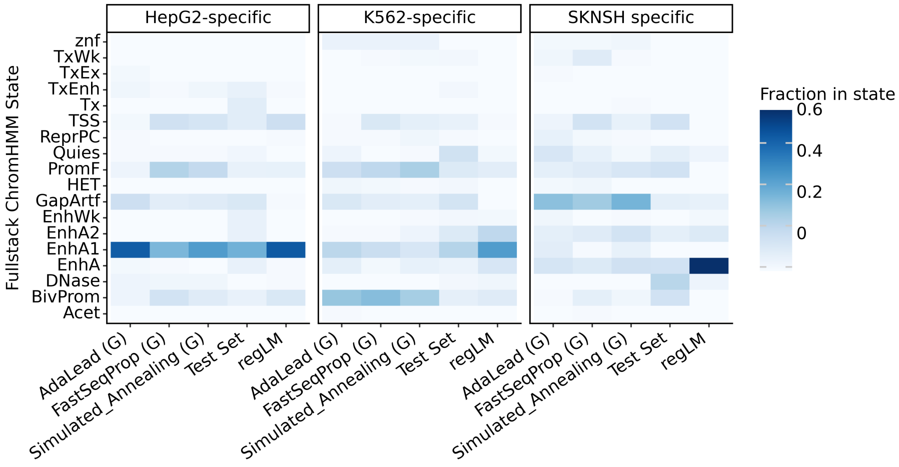


**Fig. S23.** Predictions of a classification model trained to classify genomic DNA into chromatin states defined by the fullstack chromHMM annotation [6], on synthetic cell type-specific enhancers generated by different methods. (G) indicates that the method was performed by Gosai et al.[4]. The 300 Gosai et al. [4] designed elements chosen based on similar activity to regLM generated enhancers are shown here.

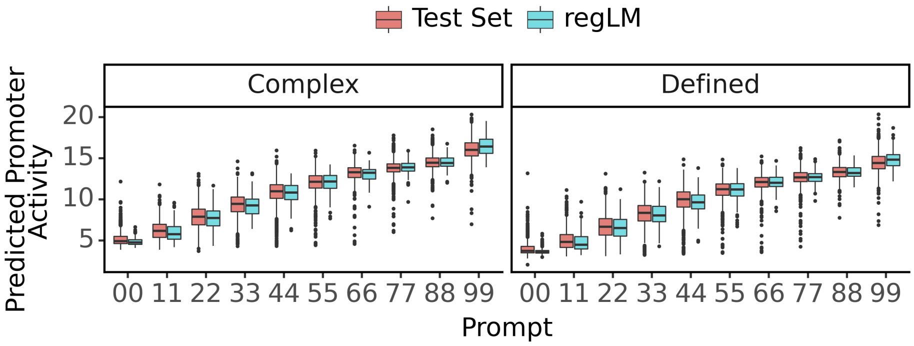


**Fig. S24.** The yeast regLM model was re-trained with labels consisting of 10 tokens ranging from 0 (lowest activity) - 9 (highest activity). The trained model was prompted with labels ranging from 00-99 and 100 synthetic promoters were generated from each prompt. The activity of these synthetic promoters was predicted using regression models trained on separate data and compared to the predicted activity of experimentally validated test set promoters with the same labels.
